# Recognition of small Tim chaperones by the mitochondrial Yme1 protease

**DOI:** 10.1101/2025.07.23.666395

**Authors:** Mariella Quispe-Carbajal, Lauren Todd, Steven E. Glynn

## Abstract

Yme1 is a conserved ATP-dependent protease that maintains mitochondrial function by degrading proteins in the intermembrane space. However, how Yme1 selects substrates within the crowded mitochondrial environment is poorly understood. An established substrate of Yme1 in yeast is the Tim10 subunit of the small Tim9-Tim10 protein chaperone complex, which is degraded following disruption of the subunit’s internal disulfide bonds. Here, we use biochemical and biophysical approaches to examine initial substrate binding and degradation of small Tim proteins by Yme1 and shed light on the molecular mechanism of substrate selection. We show that Yme1 preferentially binds Tim10 over other small Tim proteins by forming a high-affinity interaction with the subunit irrespective of the presence of its disulfide bonds. This interaction is primarily mediated by Tim10’s flexible N-terminal ‘tentacle’, though substrate unfolding exposes additional contact sites that enhance engagement. Notably, the human ortholog TIMM13 is also recognized by yeast Yme1, suggesting conservation of recognition strategy across species. Yme1 also binds to the assembled Tim9-Tim10 chaperone but independently of the Tim10 N-terminal tentacle. These findings suggest that Yme1 surveils the folding state of Tim10 throughout its functional lifecycle - both as a folded monomer and as a subunit of the functional chaperone complex - but only commits to degradation after disruption of its disulfide bonds.

## Introduction

Mitochondria use sophisticated protein quality control systems to maintain their function[1, 2]. Imported polypeptides must be properly folded and sorted to their correct compartment; and damaged, unassembled and unnecessary mitochondrial proteins must be removed to prevent aggregation and maintain proper stoichiometry of large complexes[2-4]. Disruption of mitochondrial quality control results in cellular stress and the development of severe human diseases[1, 5, 6]. A major component of this quality control system is the highly conserved ATP-dependent protease Yme1 (YME1L in humans) [7-9]. Individual Yme1 subunits contain a small N-terminal domain, a single-pass transmembrane helix, a AAA+ ATPase domain and a C-terminal zinc metalloprotease domain[8]. These subunits assemble into active hexameric proteases that are inserted in the inner membrane, with an enzymatic core of ATPase domains sitting atop a proteolytic chamber that projects into the intermembrane space (IMS). From this position, Yme1 can access a diverse collection of protein substrates in both the inner membrane and IMS and translocate them into the chamber for degradation[7, 10, 11]. The question of how Yme1 selects these substrates for irreversible proteolysis from the crowded mitochondrial environment lies at the heart of its role in quality control.

A well-established substrate of yeast Yme1 is Tim10, a member of the highly conserved small Tim family of mitochondrial chaperones that escort imported hydrophobic membrane proteins across the IMS prior to insertion into either the outer or inner membranes[12]. These small Tim proteins assemble into distinct ring-like hexamers, each containing alternating copies of two subunits (known as Tim8-Tim13 and Tim9-Tim10 in yeast; TIMM8a-TIMM13 and TIMM9-TIMM10 in humans)[13]. Tim9-Tim10 is essential for viability in yeast and helps chaperone key membrane proteins such as the ADP/ATP carrier [14-16]. All small Tim subunits share a simple helix-turn-helix structure with disordered ‘tentacles’ of varying lengths extending from their N- and C-termini[17, 18]. The helices are spanned by twin disulfide bonds encoded by Cys-X_3_-Cys motifs and formed during import into the mitochondria by the Mia40-Erv1 oxidative folding system[19]. In the yeast Tim9-Tim10 complex, disruption of these disulfides prompts accumulation of unassembled monomers and degradation of Tim10 subunits by the Yme1 protease [20, 21].

Previous biochemical studies have shown that Yme1 exhibits a strong preference for degrading Tim10 compared to Tim9, despite their high sequence similarity, and that chemical reduction of the Tim10 disulfide bonds prompts unfolding of the subunit and rapid degradation *in vitro* [22].

This degradation is mediated by the N-terminal tentacle of Tim10, which can act as an autonomous degron sequence when attached to unrelated protein domains [22]. However, the precise nature of the interaction between Yme1 and its substrates remains poorly understood. Substrate-engaged cryo-EM structures of Yme1 and the related human AFG3L2 protease revealed how pore loops in the ATPase domains grip polypeptides as they are translocated into the proteolytic chamber. However, these structures likely captured an advanced stage in the degradation cycle, and not than the initial substrate-binding event that determines specificity [23, 24]. Co-immunoprecipitation experiments in yeast identified two distinct helical regions on the Yme1 surface that bind differentially to the inner membrane proteins Cox2 and Phb1, supporting the concept of substrate-specific binding sites on the enzyme surface [25].

How Yme1 chooses specific substrates from the IMS and discriminates between functional and destabilized proteins for degradation is a key question in mitochondrial protein quality control. Here, we investigate the molecular basis of Yme1 substrate recognition by examining the initial binding event between the protease and small Tim proteins. We show that degradation of a preferred substrate, Tim10, is driven by a high affinity interaction that does not require the protease to adopt a specific nucleotide state and is formed irrespective of the presence of disulfide bonds within the substrate. While this binding is primarily mediated by the Tim10 N-terminal tentacle, substrate unfolding exposes additional contact sites that enhance engagement by the protease. Comparative analyses of interactions between Yme1 and the yeast and human small Tim proteins suggest that some human chaperones may employ a similar mode of recognition. Finally, we show that Yme1 also binds the assembled Tim9-Tim10 chaperone complex in a manner independent of the Tim10 tentacle, suggesting that Yme1 closely monitors the folding state of Tim10 subunits throughout their functional lifecycle and only commits to degradation following disruption of their disulfide bonds.

## Results

### Yme1 binds to Tim10 with high affinity independent of its disulfide bonds

We previously generated rebuilt Yme1 proteases where the insoluble transmembrane domains are replaced by a soluble hexamerization domain (cc-hex) that drives assembly of enzymatically active hexamers (hexYme1) [22, 26]. Using these rebuilt proteases, we developed a microscale thermophoresis (MST) assay that quantifies protein substrate affinity by measuring the thermal motion of fluorescently labeled proteases in response to binding **(Fig. 1A)**. NTA-fluorescent dye labels were added to a hexYme1 construct bearing a C-terminal His_6_ tag (hexYme1^His^), which has comparable degradation activity to the untagged enzyme **(Fig. 1A)**. Labeled proteases were incubated with putative substrate proteins in the presence of the non-hydrolysable ATP analogue, ATPγS, which locks the protease in an ‘ATP-state’ and blocks substrate translocation to allow capture of initial binding events. Measurement of dissociation constants between hexYme1^His^ and oxidized Tim10 or Tim9 monomers revealed a high affinity interaction with Tim10 (Kd = 2.0 ± 0.3 μM) that is absent for Tim9 (60.0 ± 9.3 μM) (**Fig. 1B and 1D**). As previously observed, both oxidized monomers were very slowly degraded by hexYme1 (Tim10 k_deg_ = 0.09 ± 0.004 min^-1^ enz_6_^-1^; Tim9 k_deg_ = 0.02 ± 0.01 min^-1^ enz_6_^-1^), whereas Tim10 (0.44 ± 0.01 min^-1^ enz_6_^-1^) but not Tim9 (0.09 ± 0.005 min^-1^ enz_6_^-1^) was rapidly degraded following chemical reduction of the disulfide bonds with DTT **(Fig. 1D; Fig. S1)[22]**. Circular dichroism and NMR spectra of reduced Tim9 and Tim10 have previously confirmed subunit unfolding under these conditions [22, 27]. Surprisingly, despite the increased degradation rate of DTT-treated Tim10, we observed only a small but statistically significant increase in binding affinity for the protein in the reduced state (1.3 ± 0.1 μM) (**Fig. 1C**). Reduction of Tim9 did produce a notable increase in affinity (25.1 ± 2.4 μM) but at a considerably higher Kd than observed for Tim10 in either state **(Fig. 1C and 1D)**. Together, these results demonstrate that the preference of Yme1 for degrading Tim10 is driven by an initial high affinity interaction between substrate and protease but that this interaction occurs irrespective of the disulfide bonds within Tim10. Furthermore, Yme1 exhibits an apparent preference for binding to unfolded small Tim proteins even without the presence of a suitable degron sequence.

**Figure 1.**
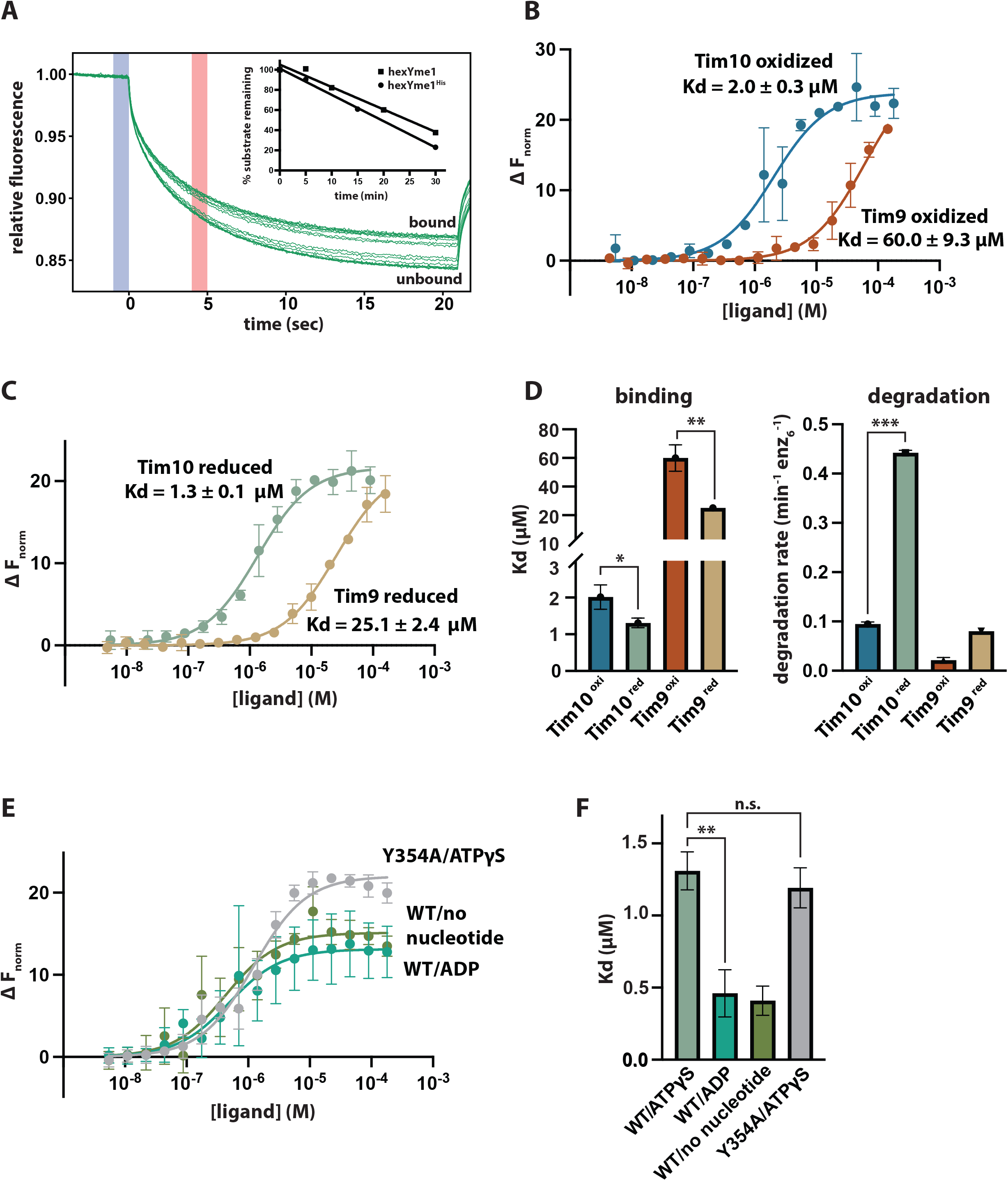
Yme1 forms a high affinity interaction with Tim10 independent of disulfide bonds or nucleotide state. (A) Representative MST traces showing response in the unbound and bound states. Samples contained 50 nM labeled hexYme1^His^, 5 mM ATPγS and between 5.5-9000 nM Tim10. Inset: degradation of reduced Tim10 (10 µM) by either C-terminally tagged hexYme1^His^ or untagged hexYme1 (B) Equilibrium binding curves showing interaction of oxidized Tim10 or oxidized Tim9 to hexYme1^His^ in the presence of 5 mM ATPγS. (C) Equilibrium binding curves showing interaction of reduced Tim10 and reduced Tim9 to hexYme1^His^ in the presence of 5 mM ATPγS. (D) Dissociation constants calculated from MST data shown in Fig. 1B and 1C. Error bars represent standard error of Kd. Initial degradation rates for oxidized and reduced Tim9 and Tim10 (10 µM) by hexYme1 (0.5 µM). (E) Equilibrium binding curves showing interaction of reduced Tim10 to hexYme1^His^ or in the presence of 5 mM ATPγS, ADP, or with no added nucleotide; and ^Y354A^hexYme1^His^ in the presence of ATPγS. (F) Dissociation constants calculated from MST data shown in Fig. 1E. All binding curves and initial degradation rates are means of independent replicates (n=3) ± s.d. *p ≤ 0.05, **p ≤ 0.01, ***p ≤ 0.001 as calculated using the Student’s two-tailed t-test.

AAA+ proteases can adopt different conformations dependent on their nucleotide state and in some cases, bind to substrates only in the presence of ATP or its analogues[23, 28]. To examine the nucleotide-dependency of substrate binding by Yme1, we measured binding to reduced Tim10 in the presence of either ADP (0.5 ± 0.2 μM) or with no added nucleotide (0.4 ± 0.1 μM) **(Fig. 1E and 1F)**. In both cases, the affinity of Yme1 for Tim10 showed a small but statistically significant increase compared to measurements with ATPγS, demonstrating that Yme1 does not require a specific nucleotide state to create the high affinity interaction with Tim10. To exclude the possibility that our measured affinities were affected by slow translocation of the substrate into the degradation chamber, rather than solely reporting on the initial interaction at the protease surface, we employed a variant of Yme1 where the pore-1 loop aromatic residues are substituted to Ala (^Y354A^hexYme1^His^). This substitution has been shown to block substrate translocation by Yme1 and other AAA+ proteases [23, 29]. We observed no significant change in affinity for binding of reduced Tim10 to this variant in the presence of ATPγS (1.2 ± 0.1 μM), indicating that these residues do not participate in substrate binding and confirming that our experimental approach is capturing the initial binding event **(Fig. 1E and 1F)**.

### The interaction with Tim10 is primarily mediated by the N-terminal tentacle

Our previous studies showed that fusing the N-terminal tentacle of Tim10 to unrelated proteins can initiate their degradation by Yme1 *in vitro* [22]. Consistent with this observation, truncation of the entire fourteen residue tentacle of Tim10 (^ΔN14^Tim10) significantly impaired binding and abolished degradation of the oxidized monomer (Kd = 45.6 ± 6.5 μM; k_deg_ = 0.01 ± 0.004 min^-1^ enz_6_^-1^) **(Fig. 2A and 2B)**. Chemical reduction of the truncated substrate partially rescued binding but not degradation (Kd = 5.9 ± 0.5 μM; k_deg_ = 0.02 ± 0.008 min^-1^ enz_6_^-1^). This suggests that unfolding of Tim10 exposes additional elements that are recognized by Yme1 but that the N-terminal tentacle is required for translocation and degradation **(Fig. 2A and 2B)**. Moreover, incorporation of two negatively charged Asp residues within the substrate N-terminus (Ser2Asp, Phe3Asp; ^DD^Tim10), which has been shown to block the degradation of many AAA+ protease substrates, also impaired binding of the oxidized monomer and greatly reduced degradation even following chemical reduction (Kd ^DD^Tim10^oxi^ = 21.2 ± 2.2 μM; k^deg DD^Tim10^red^ = 0.03 ± 0.03 min^-1^ enz_6_^-1^), further supporting the importance of the N-terminal tentacle in mediating the initial interaction **(Fig. 2A and 2B)**.

**Figure 2.**
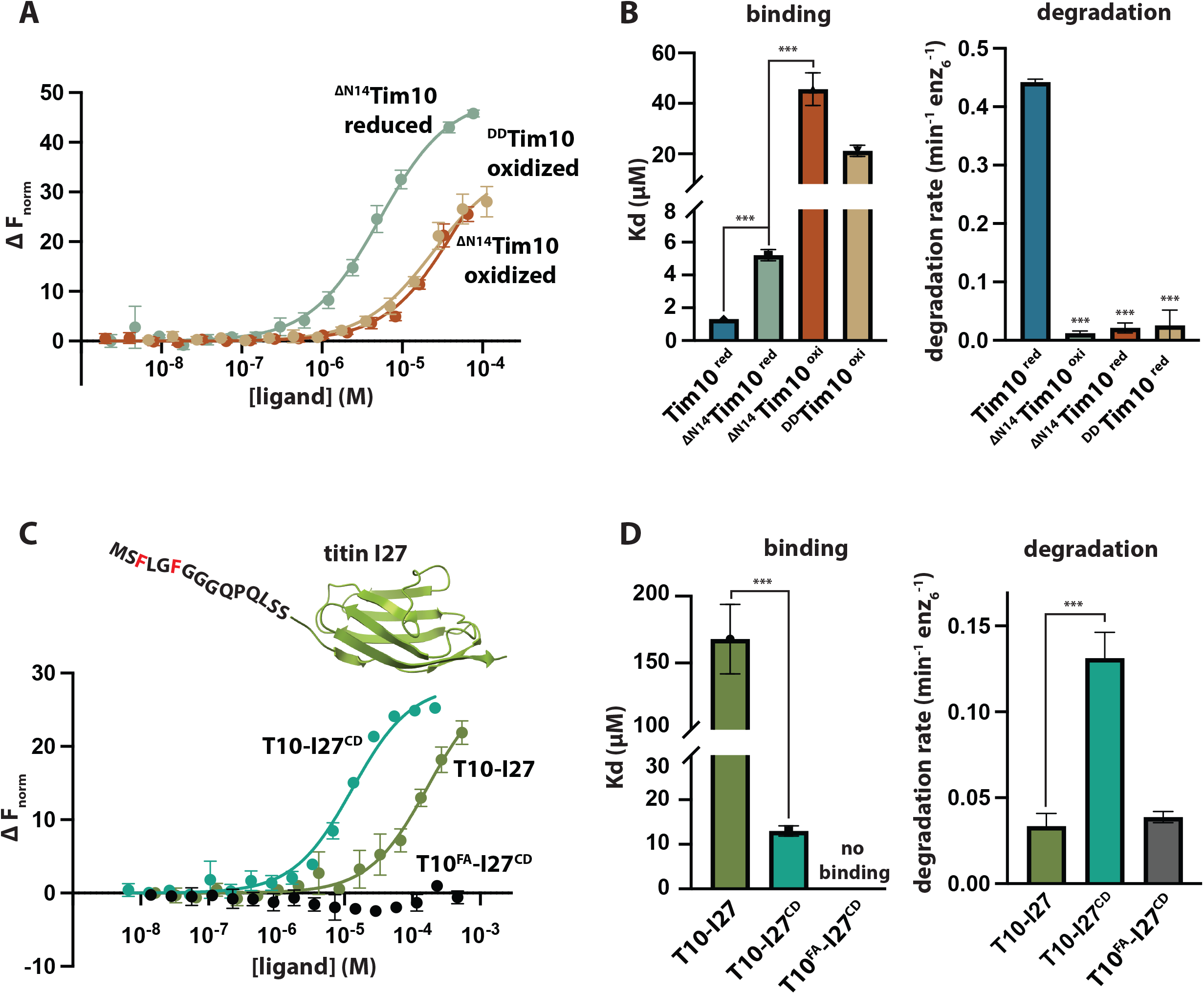
The interaction of Tim10 with Yme1 is largely mediated by the flexible N-terminus. (A) Equilibrium binding curves of oxidized and reduced ^ΔN14^Tim10 and oxidized ^DD^Tim10 to hexYme1^His^ (B) Dissociation constants calculated from MST data shown in Fig. 2A. Error bars represent standard error of Kd. Initial degradation rates for oxidized and reduced ^ΔN14^Tim10 and reduced ^DD^Tim10 (10 µM) by hexYme1 (0.5 µM). (C) Equilibrium binding curves of T10-I27, T10-I27^CD^ and T10^FA^-I27^CD^ to hexYme1^His^. (D) Dissociation constants calculated from MST data shown in Fig. 2C. Error bars represent standard error of Kd. Initial degradation rates for I27 variants (10 µM) by hexYme1 (0.5 µM). All binding measurements were performed in the presence of 5 mM ATPγS. All binding curves and initial degradation rates are means of independent replicates (n=3) ± s.d. ***p ≤ 0.001 as calculated using the Student’s two-tailed t-test.

To investigate if the N-terminal tentacle alone is sufficient to promote interaction with the protease, we attached residues 1-14 of Tim10 to both the folded I27 domain of human titin (T10-I27) and a soluble unfolded variant generated by substitution of two internal Cys residues to Asp (T10-I27^CD^). Surprisingly, folded T10-I27 exhibited very weak binding (Kd >150 μM) and a very slow rate of degradation (0.03 ± 0.01 min^-1^ enz_6_^-1^), whereas unfolded T10-I27^CD^ did bind (13.0 ± 1.2 μM), albeit with a weaker affinity than full-length Tim10, and was degraded more rapidly than its folded counterpart (0.13 ± 0.02 min^-1^ enz_6_^-1^) **(Fig. 2C and 2D)**. Within the Tim10 N-tentacle, residues Phe-3 and Phe-5 have been suggested to contribute significantly to degradation [22]. Substitution of these two residues to Ala in full-length Tim10 resulted in protein aggregation and prevented quantification of binding. However, an unfolded T10-I27^CD^ construct bearing these same two substitutions (T10^FA^-I27^CD^) was soluble and displayed both complete loss of binding to Yme1 and very slow degradation (0.04 ± 0.003 min^-1^ enz_6_^-1^) **(Fig. 2C and 2D)**.

Together, these results suggest that: (1) the Tim10 N-terminal tentacle is primarily responsible for the initial binding event and that one or both of residues Phe-3 and Phe-5 are required for this interaction and (2) Yme1 prefers to bind to suitable degron sequences when they are part of an unstructured polypeptide.

### Cross-species recognition of small tim proteins

Yeast contain additional small Tim proteins in the IMS and orthologues of these subunits are conserved across eukaryotes. An analysis of AlphaFold3 predicted structures of small Tim proteins from both *S. cerevisiae* and humans revealed that all proteins, with the exception of human TIMM10, contain flexible N-terminal tentacles of at least 6 residues **(Fig. 3A; Fig. S2)**[30]. Of these N-terminal sequences, only yeast Tim10 and human TIMM13 contain multiple Phe residues. To ask if the human small Tim subunits contain similar recognition elements to the yeast proteins, we measured the degradation of reduced human TIMM8A, TIMM10 and TIMM13 by yeast hexYme1 (TIMM9 was excluded due to poor solubility as a monomer). Of these three human subunits, only TIMM13 was significantly degraded and notably, at a faster rate than observed for Tim10 (k_deg_ = 0.66 ± 0.33 min^-1^ enz_6_^-1^) **(Fig. 3B)**. As anticipated based on its observed degradation, TIMM13 bound to hexYme1 but with only moderate affinity (Kd = 20.0 ± 10.1 μM) compared to yeast Tim10. Importantly, substitution of the two Phe residues within the N-terminal tentacle of human TIMM13 to Ala (Phe5Ala, Phe9Ala; TIMM13^FA^) abolished binding, indicating a similar mode of recognition for the two proteins **(Fig. 3C)**.

**Figure 3.**
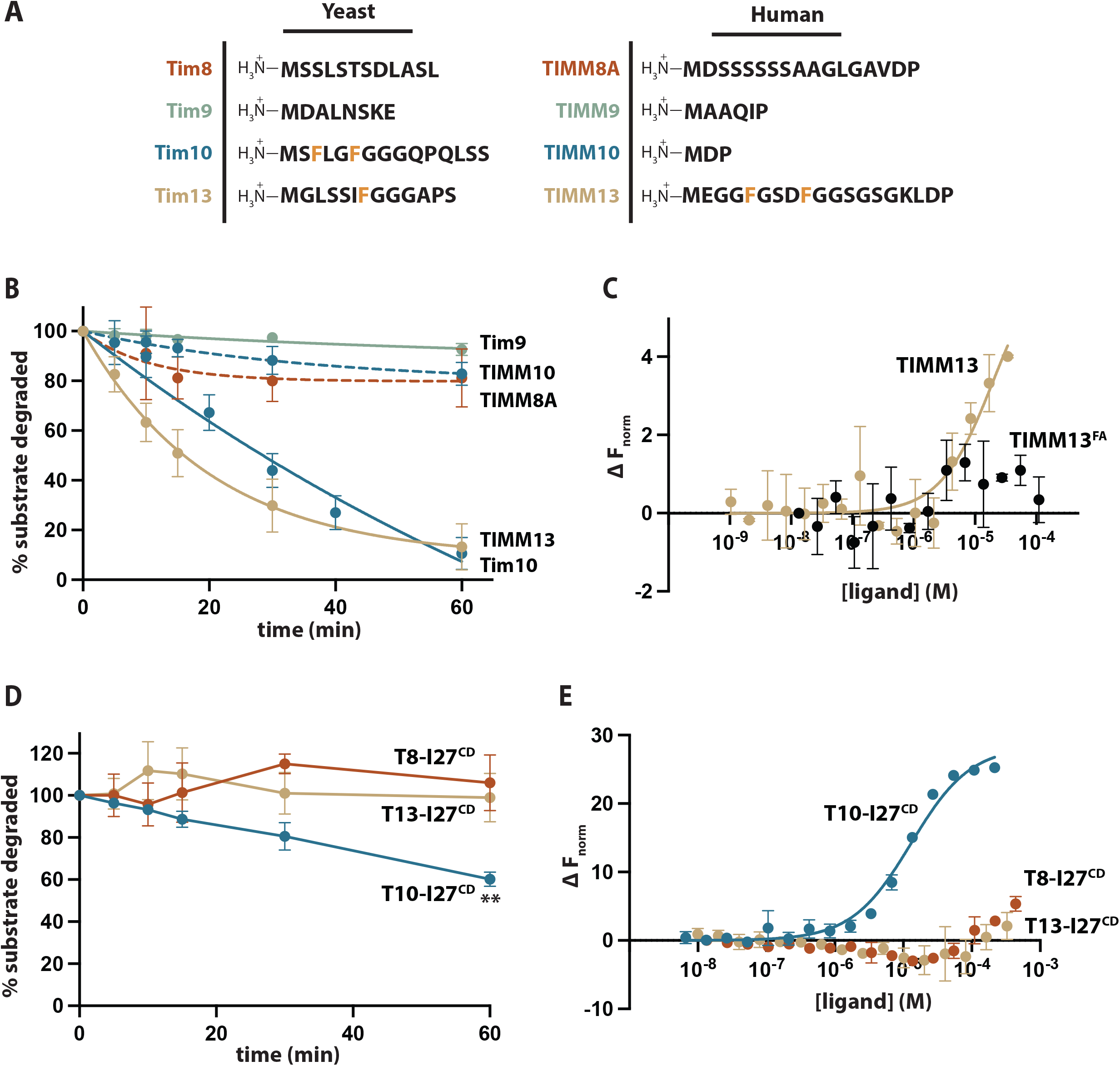
Substrate recognition small Tim proteins from human and yeast. (A) Sequences of small Tim protein N-terminal tentacles from yeast and humans. Lengths of unstructured tentacle regions were defined from AlphaFold3 predictions. (B) Plot showing degradation of chemically reduced small Tim proteins by hexYme1. Both yeast Tim10 and human TIMM13 are robustly degraded by the protease whereas yeast Tim9, and human TIMM10 and TIMM8A are not significantly degraded. (C) Equilibrium binding curves for human TIMM13 and the TIMM13^FA^ variant with hexYme1^His^. (D) Plot showing degradation of proteins containing the yeast Tim8, Tim10, and Tim13 N-terminal tentacles fused to unfolded I27^CD^ (T8-I27^CD^, T10-I27^CD^ and T13-I27^CD^). (D) Equilibrium binding curves for T8-I27^CD^, T10-I27^CD^ and T13-I27^CD^ binding to hexYme1^His^. All binding measurements were performed in the presence of 5 mM ATPγS. All binding curves and initial degradation rates are means of independent replicates (n=3) ± s.d. **p ≤ 0.01 as calculated using the Student’s two-tailed t-test.

Analysis of the remaining small Tim proteins in yeast was limited by the low solubility of full-length Tim13. To address this, we fused the N-terminal tentacles of both Tim8 (residues 1-16) and Tim13 (1-13) to unfolded I27^CD^ (T8-I27^CD^; T13-I27^CD^). In contrast to the Tim10 tentacle, which can act as an autonomous degron when fused to unrelated proteins, these other sequences did not support any appreciable binding or degradation by the protease **(Fig. 3D and 3E)**. These results suggest that Tim10 is the only small Tim protein in yeast that is recognized by Yme1 through its N-terminal tentacle and that the presence of multiple Phe residues within a flexible sequence is predictive of recognition by Yme1. Furthermore, the cross-recognition of human TIMM13 by yeast Yme1 opens the possibility that the human and yeast proteases employ similar strategies for selecting substrates.

### Yme1 binds to the Tim9-Tim10 chaperone complex independently of the N-terminal tentacle

In yeast cells, formation of hexameric Tim9-Tim10 chaperones protects the individual subunits from degradation by Yme1 [20, 21]. The tentacles of both Tim9 and Tim10 subunits are disordered in the crystal structure of the assembled chaperone, but the positions of their flanking regions suggest they extend away from the body of the heterohexamer, where they could in principle still interact with the protease **(Fig. 4A)**. To test this possibility, we assembled Tim9-Tim10 complexes by mixing the purified individual subunits at room temperature and isolating single peak representing heterohexamers by size exclusion chromatography **(Fig. 4B)**[18]. In line with *in vivo* experiments, Tim10 subunits in these complexes were resistant to degradation by Yme1 but were degraded following incubation with 10 mM DTT and 2 M urea to promote subunit reduction and disassembly of the complex **(Fig. 4C)[20, 21].** We observed a moderate affinity interaction between the assembled complex and Yme1 in our MST assay (Kd = 11.5 ± 1.6 μM) **(Fig. 4D and 4E)**. To assess whether this binding is mediated by the N-terminal tentacles of the three Tim10 subunits in the complex, we assembled variant heterohexamers containing Tim10 subunits bearing an N-terminal truncation. Although incorporation of the previously employed ^ΔN14^Tim10 subunits resulted in unstable complexes, an alternative variant missing only the first seven residues (^ΔN7^Tim10) successfully produced stable heterohexamers with Tim9 **(Fig. 4B)**. This truncated subunit lacks the key Phe residues in the tentacle and did not bind to Yme1 as an isolated monomer **(Fig. S3A)**. Surprisingly, incorporation of these ^ΔN7^Tim10 subunits did not significantly impact the binding affinity (Tim9-^ΔN7^Tim10; 14.7 ± 2.5 μM), demonstrating that Yme1 binding to the chaperone complex does not require the degron sequence **(Fig. 4D and 4E)**.

**Figure 4.**
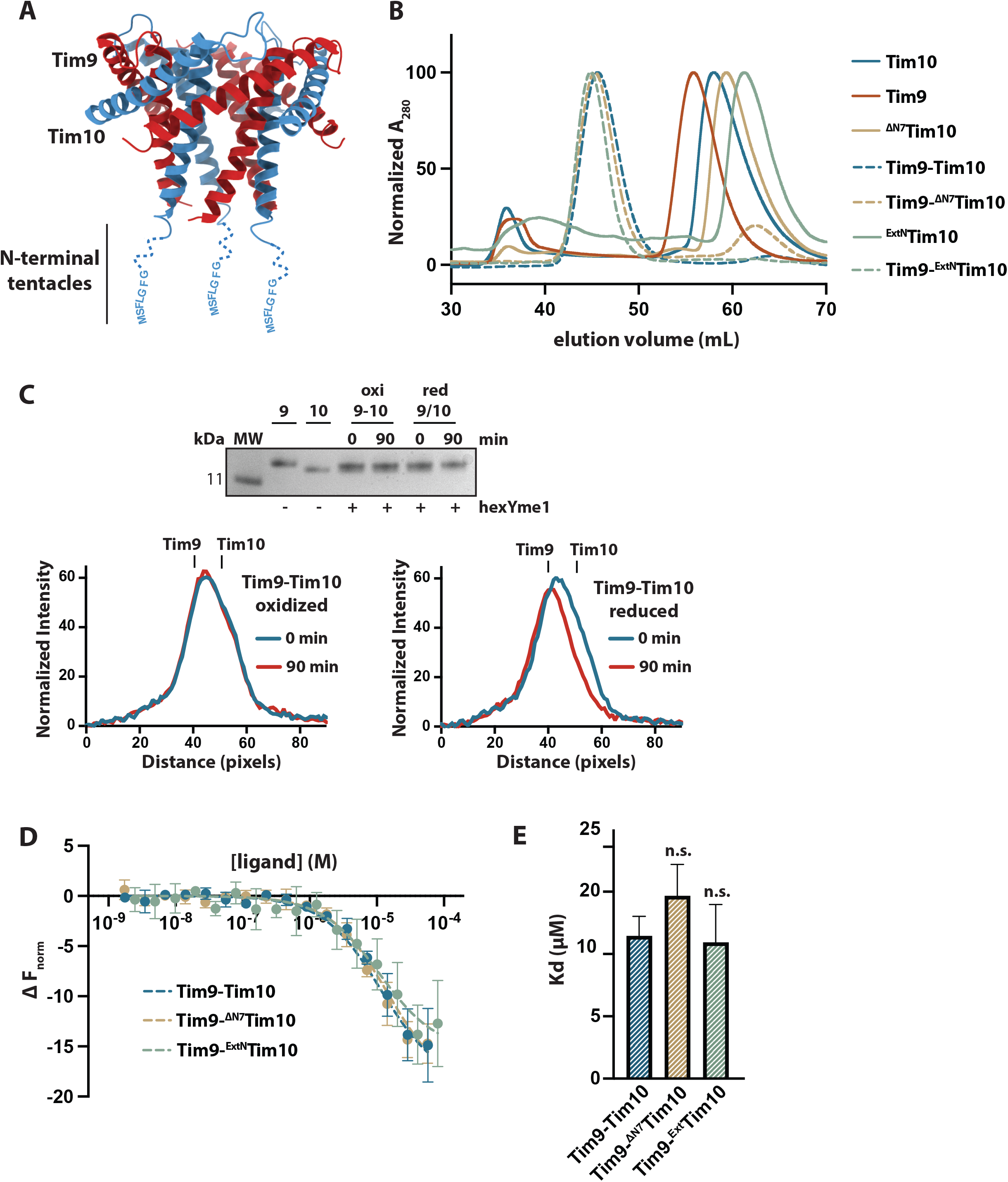
Yme1 binds to Tim9-Tim10 complexes independently of Tim10 N-terminal tentacles. (A) Crystal structure of the *S. cerevisiae* Tim9-Tim10 chaperone complex (PDB ID: 3DXR). N-terminal tentacles have been added for illustrative purposes. (B) Superdex 100 10/300 GL size exclusion chromatography profiles of isolated Tim 10 and Tim 9 subunits, wild type Tim9-Tim10 assembled complexes, and mutant complexes containing Tim10 variants. (C) The assembled Tim9-Tim10 complex is resistant to degradation by Yme1. SDS-PAGE showing migration of isolated Tim9 (9) and Tim10 (10) monomers and doublet bands corresponding to subunits of the assembled Tim9-Tim10 complex following 90 min incubation with hexYme1. No change in band intensity for the doublet is seen for oxidized Tim9-Tim10 complex with hexYme1. Incubation of DTT-treated Tim9-Tim10 complex with hexYme1 results in loss of lower band in the doublet. Band intensity profiles are also shown for both oxidized (left) and DTT-treated Tim9-Tim10 (right) doublets before (blue) and after (red) incubation with hexYme1 as quantified in ImageJ. Migration positions of monomeric Tim9 and Tim10 are shown. The band profile for only the DTT-treated Tim9-Tim10 complex shows loss of intensity at the position corresponding to Tim10 following incubation with hexYme1, indicating selective degradation of that subunit. (D) Equilibrium binding curves of hexYme1^His^ binding to the Tim9-Tim10 wild-type complex and variant complexes containing either N-terminally truncated Tim10 subunits (Tim9-^ΔN7^Tim10) or Tim10 subunits containing an extended linker sequence between residues 15 and 16 (Tim9-^ExtN^Tim10). (E) Dissociation constants calculated from MST data shown in Fig. 4D. Error bars represent standard error of Kd. All binding measurements were performed in the presence of 5 mM ATPγS. All binding curves and initial degradation rates are means of independent replicates (n=3) ± s.d.

The observation that the Tim10 N-terminus is dispensable for binding of the complex suggests that these tentacles can no longer access their binding site on the protease when part of the assembled chaperone. This inaccessibility could result from steric hindrance imposed by the larger molecular weight chaperone complex compared to the monomer. To alleviate any potential steric constraints, we constructed a Tim10 variant containing a ten-residue flexible linker sequence (GGGSQGSGGG) located between residues 15-16 of wild-type Tim10, effectively separating the N-terminal tentacle from the beginning of the first α-helix (^ExtN^Tim10). As a monomer, ^ExtN^Tim10 bound to Yme1 with low but detectable affinity (>50 µM) **(Fig. S3C)**. However, when incorporated into Tim9-Tim10 chaperones (Tim9-^ExtN^Tim10), this complex bound to Yme1 with an affinity indistinguishable from either wild-type or the truncated Tim9-^ΔN7^Tim10 variant (10.9 ± 3.0 µM) **(Fig. 4D and 4E)**. Together, these findings indicate that Yme1 binds to the assembled chaperone in a manner that is independent of the Tim10 N-terminal tentacles.

## Discussion

Disruption of the disulfide bonds in Tim10 initiates its rapid degradation by Yme1 in both yeast cells and solution experiments [20, 22]. In our experiments, Yme1 binds with high affinity to Tim10 in the disulfide-bonded state and with less than a two-fold decrease in Kd following reduction of the disulfides and associated unfolding. Thus, the barrier that protects intact Tim10 monomers from degradation is not substrate binding but rather the failure of Yme1 to unfold and/or translocate covalently linked polypeptide chains. This is distinct from more powerful ATP-dependent unfoldases like bacterial ClpXP, which can translocate multiple cross-linked protein chains simultaneously and instead is reminiscent of the partial proteolysis of the mitochondrial ribosomal subunit, MrpL32, performed by the related m-AAA protease, which also fails to processively unfold and degrade the substrate when it encounters a highly stable structural motif [31, 32].

Our findings indicate that the N-terminal tentacle of Tim10 is primarily responsible for driving substrate interaction with Yme1, and attaching this sequence to an unrelated protein is sufficient to initiate binding. However, these experiments also imply the existence of additional contacts between the substrate and protease not mediated by the N-terminal tentacle. This is seen most strikingly in the ten-fold difference in affinity for Yme1 binding to full-length Tim10 compared to unfolded T10-I27^CD^, which contains only the first fourteen residues from the natural substrate. Moreover, reduction of a truncated Tim10 construct lacking the N-terminus partially rescued binding to Yme1, and reduction of wild-type Tim9 also produced a notable increase in affinity, albeit much weaker than observed for full-length Tim10. In these cases, disruption of the disulfide bonds and associated protein unfolding led to increased binding affinity to the protease entirely independent of the degron sequence. One explanation is that in addition to forming a higher affinity interaction with the Tim10 N-terminus, Yme1 contains secondary substrate binding sites that broadly recognize segments of unfolded small Tim subunits. The cumulative effect of multiple weaker contacts could produce a high avidity interaction that helps tether the substrate to the protease and create a preference for degrading unfolded proteins. This principle has been demonstrated repeatedly in the weak multivalent interactions of disordered proteins containing <u>S</u>hort <u>L</u>inear <u>M</u>otifs (SLiMs)[33]. Importantly, truncated Tim10 constructs lacking the N-terminal tentacle were not degraded by the protease despite exhibiting moderate affinity binding in their reduced state. Thus, it appears that Yme1 requires the Tim10 tentacle to efficiently engage the substrate with its translocation machinery and proceed with degradation.

The ability of Yme1 to bind a protein domain fused to a suitable degron sequence is dependent on the nature of the domain. Wild-type Tim10 can tightly interact with the protease in both folded and unfolded states when the N-terminal tentacle is present. However, when this same sequence is attached to the I27 domain of human titin, which has a similar molecular weight (∼10 kDa), the protease can only bind to the protein in the unfolded state. In a recent solution NMR study, Weinhaupl and co-workers demonstrated that oxidized Tim10 monomers adopt a partially molten globule state that is distinct from the ordered helical structure seen in protomers of the Tim9-Tim10 complex [27]. This molten globule state contrasts with the thermodynamically stable and rigid I27 domain and may allow oxidized Tim10 to explore a wider variety of conformations that facilitate binding[34]. It should be noted that in both our engineered proteases and in the wild-type enzyme, long linker sequences extend from the ATPase domain to either the cc-hex coiled-coil or the native transmembrane domain. Substrates accessing the central translocating pore for degradation must first navigate between these linkers, creating the possibility of either a steric or electrostatic filter that permits some protein domains to enter but blocks others.

Steric constraints may also be in play when Yme1 binds with moderate affinity to the assembled Tim9-Tim10 complex. We originally anticipated that this binding would be mediated by the tentacles of the three Tim10 subunits present in each complex, which appear to extend away from the body of the chaperone [17, 27]. However, these tentacles were dispensable for chaperone binding and affinity was not increased by adding additional residues to separate the tentacle from the chaperone ring. While these results rule out that the binding we see in MST assays of the complex are due to Yme1 interacting with a small sub-population of monomers within the sample, they also demonstrate that the interaction with the assembled chaperone is distinct from that with the Tim10 monomer. One possibility is that the greater size of the hexameric chaperone ring acts as a steric block, preventing access of the tentacles to their binding site on the protease surface. Alternatively, solution NMR measurements of the assembled complex revealed an increased rigidification of the Tim10 tentacle upon complex formation that could also interfere with the higher affinity interaction[27].

The location of the Tim10 binding site on Yme1 is currently unknown. From our studies, we can state that the conserved Tyr residues within the translocating pore-1 loops do not contribute to the initial interaction, and that the binding site is not created by a specific pattern of bound nucleotides in the ATPase domains. Indeed, the nucleotide-independent binding of Tim10 that we observe points to the possibility of non-proteolytic substrate interactions in line with *in vivo* evidence indicating a chaperone-like function of Yme1 that can sense the folding state of IMS proteins without committing to degradation [35]. Experiments examining differences in substrate profiles for Yme1 from *S. cerevisia*e and *S. pombe* identified distinct helical regions on the surface of ATPase and protease rings (termed the NH and CH regions, respectively) that bind to specific substrates [25]. Mapping these regions to the now-available cryo-EM structure of Yme1 shows that the NH region is heavily negatively charged and unlikely to interact with the largely hydrophobic Tim10 tentacle [23] **(Fig. S4)**. In contrast, the CH region contains a patchwork of charged and neutral surfaces, which could accommodate unfolded polypeptides but a mechanism for how substrates binding at the exterior of the protease chamber could be engaged by the distant translocating pore loop in the ATPase domain remains elusive.

Overall, our results suggest a model where Yme1 monitors the folding state of Tim10 throughout its functional lifecycle. The protease binds with high affinity to both properly folded and destabilized monomers, and with moderate affinity to the assembled chaperone complex. This would allow Yme1 to repeatedly catch and release Tim10 but only proceed with degradation following disruption of the disulfide bonds **(Fig. 5)**. The question of why Yme1 chooses to rapidly degrade destabilized Tim10 but not its obligate binding partner, Tim9, is intriguing. Quantification of protein abundances in yeast cells reveal that the steady-state levels of the two subunits are approximately equal despite a considerably shorter half-life for Tim10 (Tim9: 353 ppm, 8.4 hr; Tim10: 300 ppm, 3.3 hr) [36, 37]. Furthermore, measurements of relative expression levels by translation-complex profile sequencing reveal that Tim10 is translated at more than seven-fold greater levels than Tim9 [38]. Thus, Tim10 appears to be continually expressed and degraded, allowing Yme1 to maintain correct stoichiometry of the functional chaperone complex and prevent accumulation of inactive monomers.

**Figure 5.**
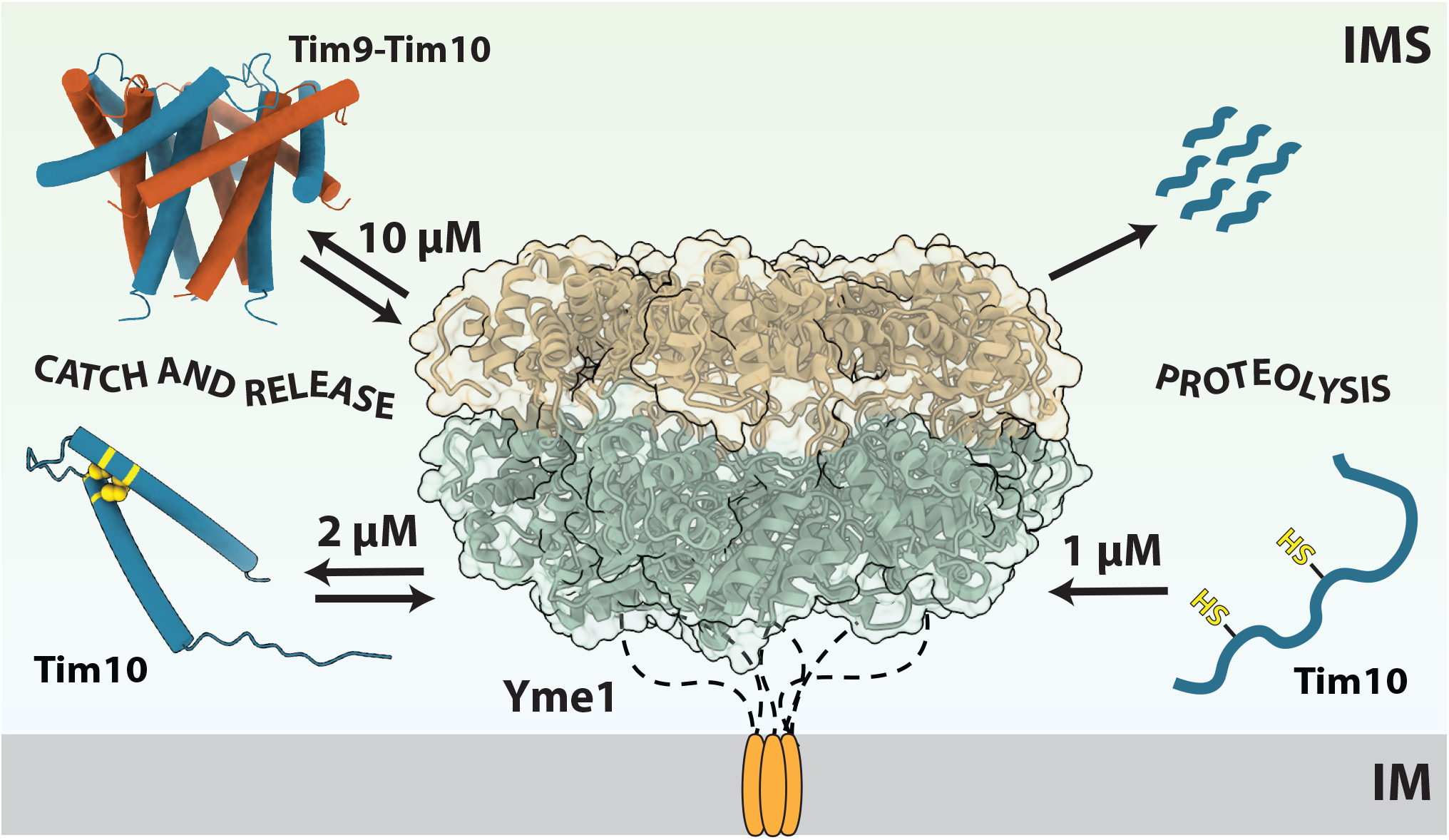
A model for surveillance of Tim10’s folding state by Yme1. Yme1 interacts repeatedly with Tim10 throughout its functional lifecycle. Moderate affinity interactions are made with the assembled Tim9-Tim10 complex and high-affinity interactions are made with isolated Tim10 monomers. Despite the high-affinity interaction with the disulfide-bonded Tim10 monomer, degradation cannot proceed unless the disulfide bonds are disrupted resulting in subunit translocation into the proteolytic chamber.

### Experimental Procedures Genes and Proteins

A plasmid containing yeast hexYme1^His^ bearing an N-terminal GST tag was generated from the existing His_6_-GST-hexYme1 expression construct by PCR amplification using primers designed to amplify the GST-hexYme1 open reading frame, followed by subcloning into the 2Bc-T vector (Addgene #37236) that encodes a C-terminal His_6_ tag [22]. Both hexYme1 and hexYme1^His^ plasmids were grown in *E. coli* BL21 CodonPlus (DE3) cells (Novagen) at 37 °C in LB supplemented with ampicillin (100 µg/mL) and chloramphenicol (34 µg/mL). Protein expression was induced by addition of 1 mM IPTG at OD_600_ ∼0.6 and grown at 16 °C for 16 h. Cells were harvested, resuspended in lysis buffer [20 mM Tris HCl pH 8.0, 300 mM NaCl, 10 mM MgCl_2_, 0.1 mM EDTA, 10 % glycerol, 10 mM 2-mercaptoethanol, 1 mM PMSF] and lysed by sonication. Clarified lysates were then applied to a gravity column containing glutathione agarose resin (Pierce). Unbound proteins were removed by washing with 20 CV lysis buffer and bound target proteins eluted with addition of 15 mL elution buffer [20 mM Tris HCl pH 8.0, 300 mM NaCl, 10 mM MgCl_2_, 0.1 mM EDTA, 10 % glycerol, 10 mM 2-mercaptoethanol, 10 mM GSH]. N-terminal GST tags were removed by addition of 1 mg TEV protease followed by incubation at 4 °C for 16 h. The post-cleavage mixture was applied to a HiLoad 26/600 Superdex 200 pg size exclusion column (Cytiva) equilibrated with gel filtration buffer [25 mM HEPES pH 8.0, 100 mM KCl, 5 mM MgCl_2_, 5 % glycerol, 1 mM DTT]. Fractions containing target protein were pooled and concentrated prior to flash freezing.

Genes encoding yeast small Tim proteins were previously cloned into the 2S-U vector that encodes an N-terminal His_6_-SUMO tag that can be removed by the Ulp1 protease to yield a scarless N-terminus [22]. Codon optimized genes encoding human small Tim proteins and fusion proteins containing human I27 or I27^CD^ domains were synthesized (Azenta) and sub-cloned into the 2S-U vector. All substrate proteins and their variants were expressed in *E. coli* Origami B pLysS (DE3) cells (Novagen) by growing cultures at 37 °C to OD_600_ ∼0.6 followed by addition of 1 mM IPTG and expression overnight at 16 °C. To purify the yeast small Tim proteins and fusion proteins containing human I27 or I27^CD^ domains, cells were harvested and resuspended in lysis buffer [20 mM Tris-HCl pH 8.0, 400 mM NaCl, 100 mM KCl, 5% glycerol, 1 mM PMSF] followed by lysis by sonication. Clarified lysates were incubated with HisPur Ni-NTA agarose resin (Thermo) for 1 hr and loaded into a gravity column. Weakly bound proteins were removed by washing with lysis buffer supplemented with 50 mM imidazole and target proteins eluted with lysis buffer supplemented with 250 mM imidazole. His_6_-SUMO tags were cleaved by overnight incubation with 0.3 mg His_6_-Ulp1 protease at 4 *°*C for 16 h. Post-cleavage mixtures were buffer exchanged into lysis buffer using an Econo-Pac 10DG desalting column (Bio-Rad) and applied to a fresh Ni-NTA agarose column to capture the His_6_-Ulp1 and His_6_-SUMO fragments. Flow-throughs containing the target proteins were collected and applied to a Sephacryl-100 10/300 GL size-exclusion column (Cytiva) pre-equilibrated with PD buffer [25 mM HEPES pH 8.0, 100 mM KCl, 5 mM MgCl_2_ and 5% glycerol]. Fractions corresponding to the target proteins were pooled, concentrated and flash frozen prior to storage at −80 °C. Human TIMM8a, TIMM10 and TIMM13 WT and mutant variants were purified as above using a modified lysis buffer [25 mM HEPES pH 7.4, 400 mM NaCl, 100 mM KCl, 5% glycerol, 1 mM PMSF] and PD buffer [25 mM HEPES pH 7.4, 100 mM KCl, 5 mM MgCl_2_ and 5% glycerol]. Human TIMM9 was purified as above using a modified lysis buffer [25 mM HEPES pH 7.8, 400 mM NaCl, 100 mM KCl, 5% glycerol, 1 mM PMSF] and PD buffer [25 mM HEPES pH 7.8, 100 mM KCl, 5 mM MgCl_2_ and 5% glycerol].

To assemble the wild-type Tim9-Tim10 chaperone complex and variants, separately purified yeast Tim9 and Tim10 subunits were mixed in a 1:1 molar ratio and incubated for 30 min at room temperature prior to loading on a Superdex 100 10/300 GL size exclusion column (Cytiva) pre-equilibrated in a buffer containing 20 mM Tris-HCl pH 8.0, 150 mM NaCl, and 5 % glycerol. Monodisperse peaks eluting at volumes corresponding to the assembled chaperone complexes (∼60 kDa) were collected and examined for the presence of both subunits by SDS-PAGE. Fractions containing approximately equal amounts of Tim9 and Tim10 subunits were pooled, concentrated and flash frozen for storage at −80°C.

### Protein degradation assays

Chemical reduction of individual small Tim proteins prior to degradation was performed using conditions that have been previously shown to produce unfolding[22]. Proteins were incubated at 30 °C in the presence of 1 mM DTT for 90 min, followed by supplementation of an equal concentration of DTT in the degradation reaction buffer. Disassembly of the Tim9-Tim10 complex prior to degradation was performed by incubation of Tim9-Tim10 (100 µM) at 60 °C in PD buffer supplemented with 2 M urea for 90 min. All degradation reactions were performed in PD buffer at 30 °C in 90 µL reactions containing 10 µM substrate, 0.5 µM hexYme1 or hexYme1^His^ and an ATP regeneration system (5 mM ATP, 16 mM creatine phosphate, 0.32 mg/ml creatine kinase). 10 µL samples were removed at appropriate time points and quenched by addition of 5 µL 4x Laemmli sample buffer and incubation at 95 °C for 5 min. Substrate degradation was visualized by SDS-PAGE using 12 % NuPage Bis-Tris mini-gels (Invitrogen) stained with Coomassie Blue R-250. Substrate band intensities were quantified using ImageJ and normalized to the creatine kinase band as a loading control[39]. Initial degradation rates were calculated from at least five time points in the linear range and values are shown as means of multiple independent replicates (n=3) ± s.d. Gel band profiles from degradation reactions of the Tim9-Tim10 complex were quantified in ImageJ using a standard 100-pixel box size that was aligned for each lane to allow comparison of the intensity against migration distance.

### Protein labeling and microscale thermophoresis

Fluorescent labeling of hexYme1^His^ was performed using the Monolith His-Tag Labeling Kit RED-tris-NTA 2^nd^ Generation (Nanotemper Technologies) according to the manufacturer’s instructions. Equal volumes of 50 nM fluorescent dye diluted in PBS-T buffer (phosphate-buffered saline supplemented with 0.05 % Tween-20) and 100 nM hexYme1^His^ diluted in PD buffer were mixed and incubated for 30 min in the dark at room temperature. Comparison of sample absorbance at 650 nm and 280 nm confirmed a degree of labeling under these conditions of ∼1.0 dye/enzyme. Binding affinities between labeled hexYme1^His^ and putative protein substrates were determined using a Monolith microscale thermophoresis instrument (Nanotemper Technologies). Firstly, a 16-step serial dilution of the substrate was performed in PD buffer supplemented with 0.05 % Tween-20 (v/v) and appropriate nucleotide (5 mM). Fluorescently labeled hexYme1^His^ was added to a final concentration of 50 nM and samples were incubated for 10 min at room temperature prior to loading into Monolith NT.115 Standard Capillaries. MST measurements were performed at 30 °C using the Red channel configuration with excitation power set to 20 % and medium MST power.

Data from MST traces and dose-response curves were extracted from MO. Affinity Analysis 2.2.7 (NanoTemper Technologies). Fluorescence responses are shown as ΔF_norm_ = F_norm, bound_ - F_norm, unbound_. Kd values were using a one-site binding equation Y =(0+(B_max_-0))*((X+[hexYme1]+Kd-SQRT((X+[hexYme1]+Kd)^2-4*X*[hexYme1]))/(2*[hexYme1])), where X is the concentration of ligand, Y is ΔF_norm_, B_max_ is the maximum binding in the same units as Y and K_d_ is the dissociation constant. The values shown in all binding curves represent the means of independent replicates (*n* = 3) ± s.d. Figures were prepared using GraphPad Prism v.10.5.0 (Graphpad Software Inc, San Diego, CA) and UCSF ChimeraX [40].

## Data Availability

Data is available upon request. Contact information

## Supporting Information

This article contains supporting information

## Acknowledgements

We thank Wali Karzai (Stony Brook University) for access to instrumentation and Jae Ho Lee (Stony Brook University) for assistance with analysis of published ribosome-profiling data.

## Funding and additional information

This study was supported by National Institutes of Health award R01GM115898.

