## Supplementary Information for "Recognition of small Tim chaperones by the mitochondrial Yme1 protease"

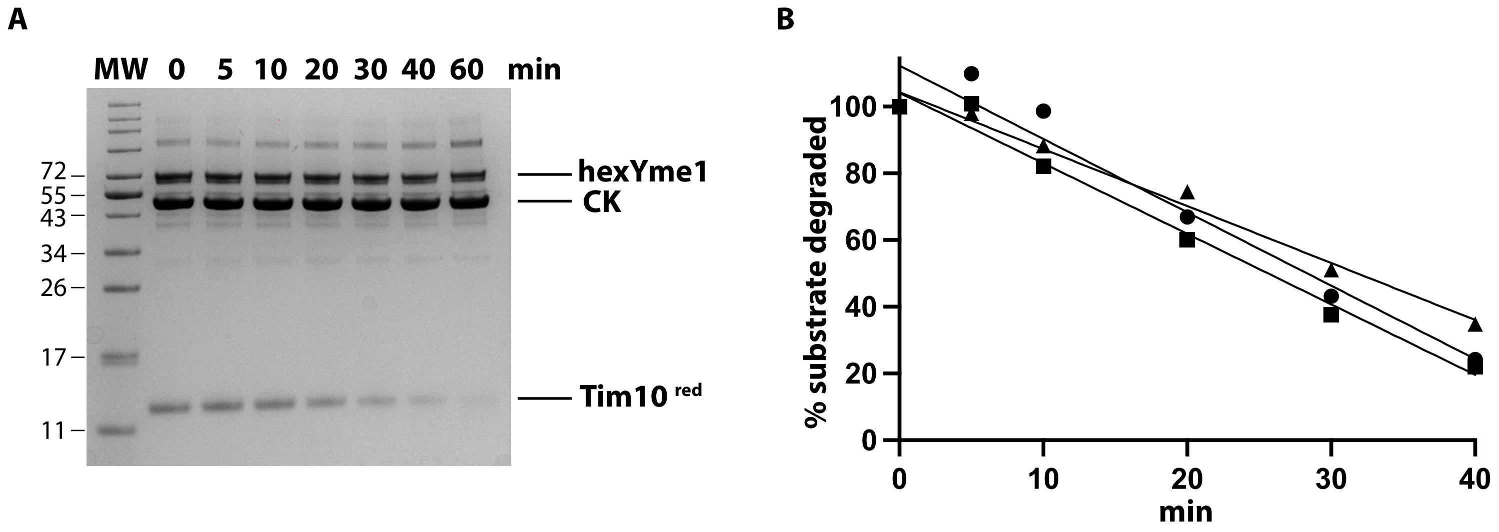


**Figure S1. Representative analysis of hexYme1 degradation. (A) SDS-PAGE showing degradation of reduced Tim10 (10 µM) by hexYme1 (0.5 µM) in the presence of an ATP regeneration system (CK). (B) Plot showing quantification of three independent replicates and linear slopes used to calculate initial rates of degradation.**

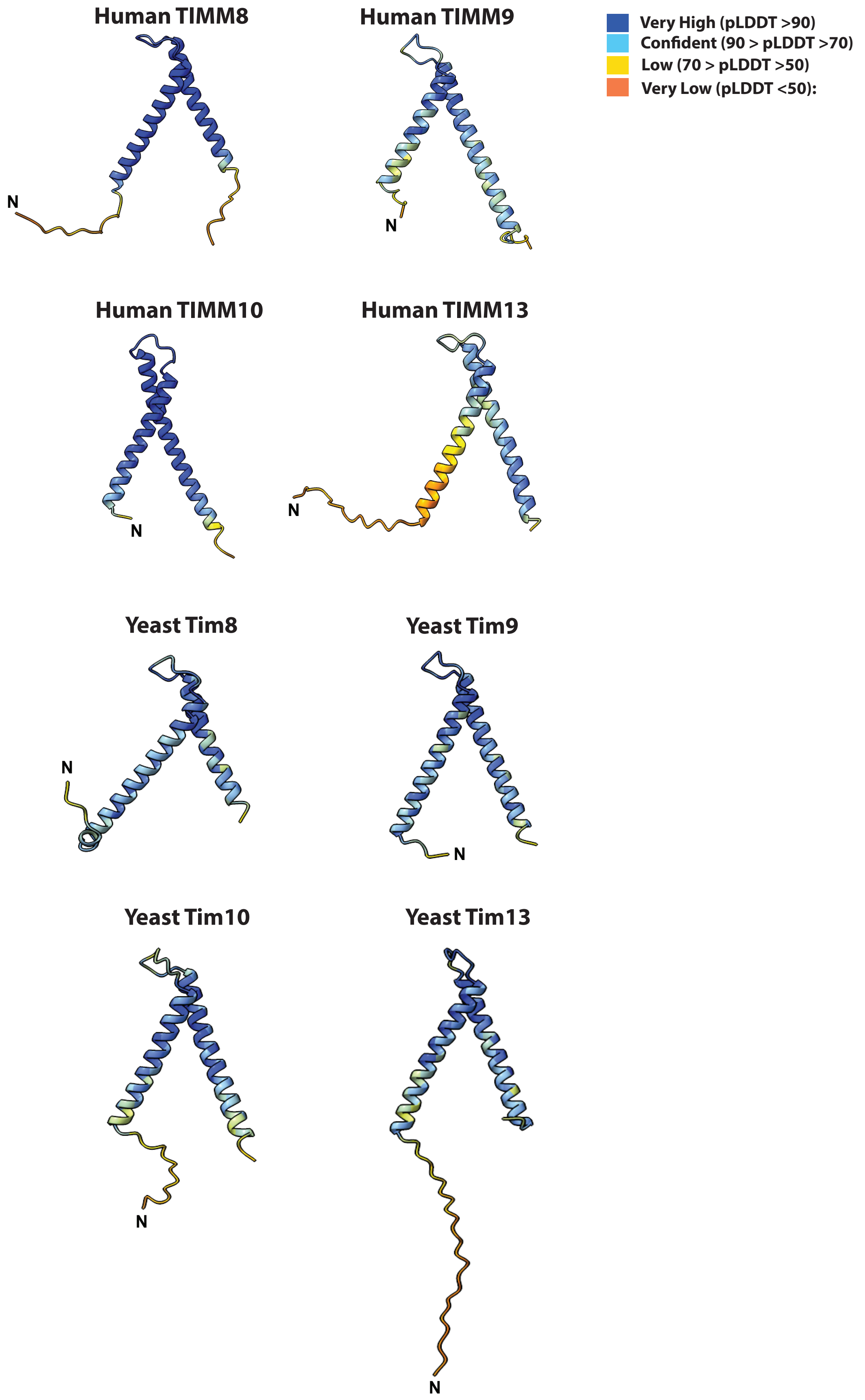


**Figure S2. AlphaFold3 predictions of small Tim proteins from human and yeast. All small Tim proteins have extended N-terminal tentacles of at least eight residues with the exception of human TIMM10. Residues are colored based on predicted local distance difference test (pLDDT).**

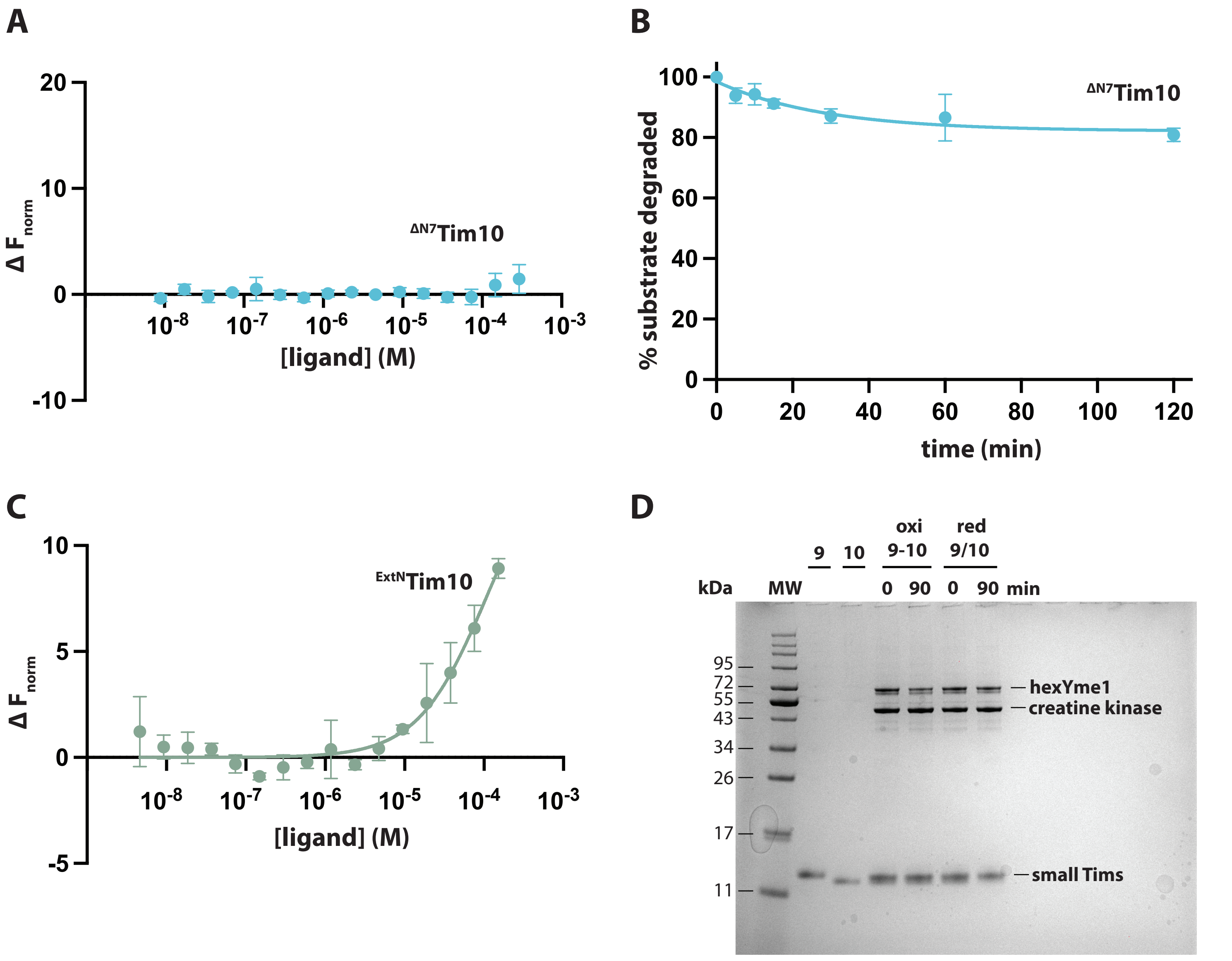


**Figure S3.** (A) Equilibrium binding data for interaction between $^{\Delta N7}$Tim10 and hexYme1$^{His}$ in the presence of 5 mM ATPγS. (B) Plot showing degradation of $^{\Delta N7}$Tim10 by hexYme1. (C) Equilibrium binding data for interaction between $^{ExtN}$Tim10 and hexYme1$^{His}$ in the presence of 5 mM ATPγS. All binding and degradation data are means of independent replicates (n=3) ± s.d. (D) Uncropped gel image from Figure 4C showing migration of Tim9 (9) and Tim10 (10) monomers alongside incubation of oxidized or reduced Tim9-Tim10 complex with hexYme1.

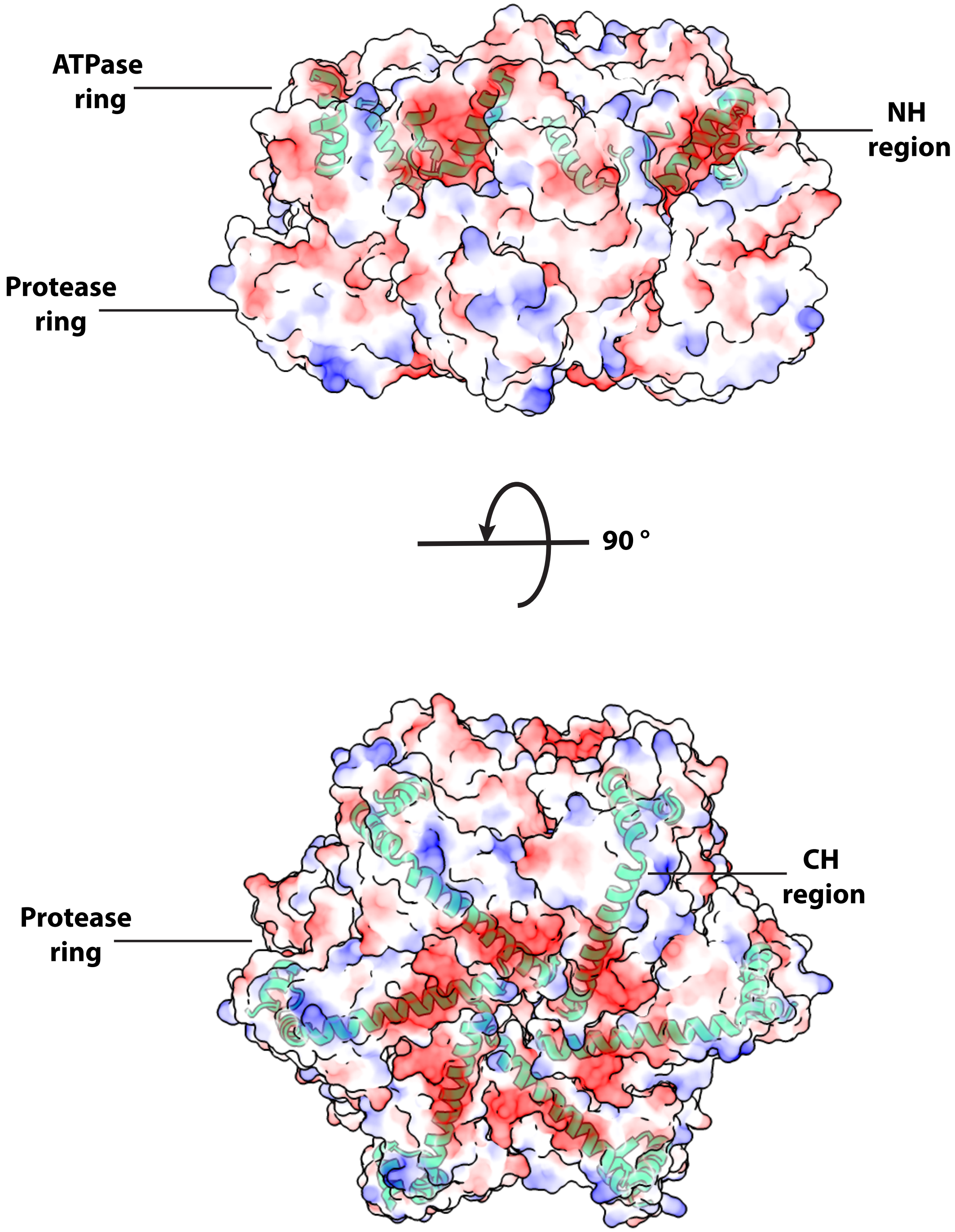


**Figure S4. Electrostatic surface representation of Yme1 (PDB ID 6AZ0) showing the location of the NH and CH regions (green helices). Images were prepared using ChimeraX and colored using the Coulombic Surface Coloring Tool**
